# Vacuolar protein sorting-associated protein 1 (Vps1) has membrane constricting and severing abilities required for endosomal protein trafficking

**DOI:** 10.1101/2025.05.25.656005

**Authors:** Shilpa Gopan, Gurmail Singh, Uma Swaminathan, Thomas J. Pucadyil

**Affiliations:** Indian Institute of Science Education and Research, Dr. Homi Bhabha Road, Pashan, Pune 411008, Maharashtra, India

**Author notes:** equal contribution (Shilpa Gopan, Gurmail Singh).

**Keywords:** Lipids, membrane templates, membrane fission, fluorescence image analysis, endosome, vacuole, protein sorting, proteomics

## Abstract

Endosomal protein trafficking pathways are fundamental to maintaining organelle identity and function and are conserved from yeast to humans. The retromer pathway traffics cargo from endosomes to the *trans* Golgi network and defects in this pathway are linked to a variety of metabolic and neurological disorders. The yeast dynamin Vacuolar protein sorting-associated protein 1 (Vps1) functions in the retromer pathway. Vps1 has been found to associate with tubular endosomes undergoing severing, implying a role in membrane fission. Current biochemical and structural analyses of Vps1 however seem to indicate that it is not sufficient for fission. Here, we analyze Vps1 functions using cell-free reconstitution, live cell imaging and proteomics of isolated yeast vacuoles. We find that the endosomal lipid phosphatidylinositol-3-phosphate (PI(3)P) stabilizes Vps1 to form protein scaffolds around tubular membrane substrates. GTP hydrolysis forces constriction of tubules to a critical ∼7 nm radius, ultimately causing their fission. Furthermore, we discover that the Insert B (InsB) in Vps1 harbor conserved lysine-rich (3K) lipid-binding and phenylalanine-rich (4F) self-assembly motifs. Mutating these motifs renders Vps1 incapable of membrane fission and abrogates its functions in endosomal protein sorting.

Quantitative vacuolar proteomic analysis reveals that loss of Vps1 results in the enrichment of a small set of proteins, most of which represent retromer cargo while causing depletion of the bulk of proteins in the vacuole. Vps1 functions in endosomal protein sorting are therefore critical for the global regulation of the vacuole proteome.

## Introduction

The intricate and finely controlled protein sorting and trafficking pathways across the endolysosomal system is fundamental to the homeostatic maintenance of organelle identity and function (Cullen, 2008). The basic design principles of these pathways appear to be evolutionarily conserved (Wendland *et al*., 1998; Burd and Cullen, 2014). Defects in these pathways have been linked to a variety of pathophysiology in humans (Lane *et al*., 2012; Sebastian *et al*., 2012; Small and Petsko, 2015; Vagnozzi and Praticò, 2019; Cullen *et al*., 2024). Trafficking pathways involve the concerted action of adaptors, coats and fission proteins to release cargo-laden transport carriers from the donor compartment. Subsequent fusion of these carriers delivers cargo to the acceptor compartment. The vacuolar protein sorting (VPS) screen in the yeast identified several proteins involved in the trafficking of cargo across the endosome or pre-vacuolar compartments (PVC) and the *trans* Golgi network (TGN) (Bankaitis *et al*., 1986; Rothman and Stevens, 1986). The Vacuolar protein sorting-associated protein 1 (Vps1) was discovered in this screen (Rothman and Stevens, 1986; Robinson *et al*., 1988). Vps1 has since been associated with the maintenance of vacuole and peroxisome morphology (Raymond *et al*., 1992; Peters *et al*., 2004; Kuravi *et al*., 2006; Vizeacoumar *et al*., 2006; Röthlisberger *et al*., 2009; Ekal *et al*., 2023), endocytosis (Nannapaneni *et al*., 2010; Rooij *et al*., 2010; Hayden *et al*., 2013; Palmer *et al*., 2015; Smaczynska-de Rooij *et al*., 2016, 2019) and autophagy (Arlt *et al*., 2023; Hu and Reggiori, 2023).

Vacuolar protein sorting involves the retromer complex that comprises of cargo-specific adaptors and membrane tubulators such as the SNX-BAR proteins. Vps1 coordinates with the retromer complex for the transport of cargo like the carboxypeptidase Y (CPY) receptor Vps10, from endosomes or the PVC to the TGN. Defects in this transport pathway result in the mis-sorting of Vps10 to the vacuole and secretion of CPY, a phenotype that formed the basis of the VPS screen (Marcusson *et al*., 1994; Nothwehr *et al*., 1995; Cooper and Stevens, 1996; Arlt *et al*., 2014; Chi *et al*., 2014). The retromer complex, comprising of the cargo-sorting trimer of Vps26, Vps29 and Vps35 along with the membrane tubulating SNX-BAR proteins, functions as a membrane coat for cargo retrieval from endosomes or the PVC (Horazdovsky *et al*., 1997; Seaman *et al*., 1998; Burda *et al*., 2002; Chi *et al*., 2014; Ma *et al*., 2017; Day *et al*., 2018; Suzuki *et al*., 2021). These coordinated activities produce cargo-laden membrane tubules, which are assumed to be severed by Vps1 (Arlt *et al*., 2014; Chi *et al*., 2014; Ma *et al*., 2017; Suzuki *et al*., 2021). Indeed, loss of Vps1 results in endosomes displaying long-lived SNX-BAR coated tubules (Chi *et al*., 2014; Ma *et al*., 2017). Furthermore, Vps1 has been found to associate with tubular membrane substrates undergoing fission (Arlt *et al*., 2014; Suzuki *et al*., 2021). Together, these observations are consistent with a model wherein Vps1 mediates membrane fission. But such a function for Vps1 has not been directly tested.

Vps1 belongs to the dynamin superfamily and contains the characteristic G domain, stalk and the bundle signalling element (BSE) (Ford and Chappie, 2019; Jimah and Hinshaw, 2019). It however contains a less characterized Insert B (InsB) in place of the membrane-binding Pleckstrin Homology Domain (PHD) found in the endocytic dynamins. InsB likely functions analogous to but is not conserved in sequence with the Variable Domain (VD) found in the mitochondrial dynamins. Dynamins self-assemble into helical scaffolds on membranes, which in turn stimulates their basal GTPase activity (Ford and Chappie, 2019). Helical scaffolds of Vps1 display an overall architecture that is more open than seen for the endocytic dynamins (Varlakhanova *et al*., 2018; Ford and Chappie, 2019). Importantly, these scaffolds lack critical interdomain contacts that are found to be necessary for stimulated GTPase activity in the endocytic dynamins. Indeed, Vps1 showed no significant stimulation in GTPase activity upon membrane binding (Varlakhanova *et al*., 2018), which is a fundamental requirement among dynamins for membrane fission (Antonny *et al*., 2016; Jimah and Hinshaw, 2019).

Here, we analyze Vps1 functions using cell-free reconstitution on templates that mimic tubulated endosomes in their lipid composition and topology, which we then correlate to endosomal sorting. Furthermore, we perform quantitative proteomic analysis of vacuoles isolated from wildtype and *vps1Δ* cells to gain insights into the repertoire of trafficking pathways requiring Vps1.

## Results

### Vps1 catalyzes fission of tubular membrane templates containing endosomal lipids

We purified a bacterially expressed Vps1 construct and tested it on membrane templates to understand its intrinsic properties. Endosomes are enriched in anionic lipids like phosphatidylserine (PS) and phosphatidylinositol 3-phosphate (PI(3)P) (Yeung *et al*., 2008; Marat and Haucke, 2016; Posor *et al*., 2022). Binding of Vps1 to these lipids was previously analyzed using vesicle sedimentation assays (Smaczynska-de Rooij *et al*., 2019), but these assays suffer from low sensitivity. Instead, we used the Proximity-Based Labeling of Membrane-Associated Proteins (PLiMAP) assay to reassess the lipid binding properties of Vps1 (Jose *et al*., 2020). PliMAP utilizes a photoactivable fluorescent lipid TDPE that crosslinks with membrane-bound proteins upon UV exposure. PliMAP assays with vesicles formed of phosphatidylcholine (PC), PC with PS (15 mol%) (PC:PS) and PC with PS (15 mol%) and PI(3)P (5 mol%) (PC:PS:PI(3)P) revealed that while Vps1 showed low binding to PC, inclusion of PS and PI(3)P significantly enhanced binding (Fig. 1A,B). Dynamins self-assemble into helical scaffolds on membranes, which in turn stimulates their basal GTPase activity (Ford and Chappie, 2019). Thus, stimulated GTPase activity reflects the fraction of membrane-bound dynamins that are self-assembled. Vps1 displayed a basal GTPase rate of ∼0.6 min^-1^ in the absence of vesicles (Fig. 1C), similar to that reported earlier (Varlakhanova *et al*., 2018). Despite residual binding to PC vesicles (Fig. 1A,B), these vesicles did not stimulate the basal GTPase rate of Vps1 (Fig. 1C), indicating that Vps1 does not self-assemble on PC vesicles. In contrast, PC:PS and PC:PS:PI(3)P vesicles stimulated the basal GTPase rate, with the latter showing a ∼ 30-fold stimulation to a rate of ∼16 min^-1^ (Fig. 1C). Thus, endosomal lipids like PS and PI(3)P promote Vps1 self-assembly and stimulate its enzymatic properties. Vps1 associates with tubular endosomes that undergo severing (Arlt *et al*., 2014; Chi *et al*., 2014; Suzuki *et al*., 2021). We therefore turned to Supported Membrane Templates (SMrTs) that display an array of pre-formed membrane tubes resting on a passive PEG cushion (Dar *et al*., 2017; Swaminathan and Pucadyil, 2024). Flowing Vps1 with GTP showed no effect on PC or PC:PS templates but templates containing PC:PS:PI(3)P underwent rapid severing (Fig. 1D, Movie S1). These results indicate that Vps1 is sufficient to cause membrane fission. To assess efficiency of the tube severing reaction, we used an image analysis routine to compute a fission index, which positively correlates with tube severing activity (see Methods). PC:PS:PI(3)P templates with GTP showed a high fission index (Fig. 1E). PC:PS or PC templates with GTP showed a fission index that was close to zero (Fig. 1E). Furthermore, PC:PS:PI(3)P templates with the non-hydrolyzable GTP analog, GppNHp or with GDP or in the absence of nucleotides also showed a fission index that was close to zero (Fig. 1E). These results signify the importance of robust membrane binding and stimulated GTPase activity for Vps1-catalyzed membrane fission. Hence, all subsequent analysis of Vps1 functions was carried out on PC:PS:PI(3)P templates.

**Fig. 1.**
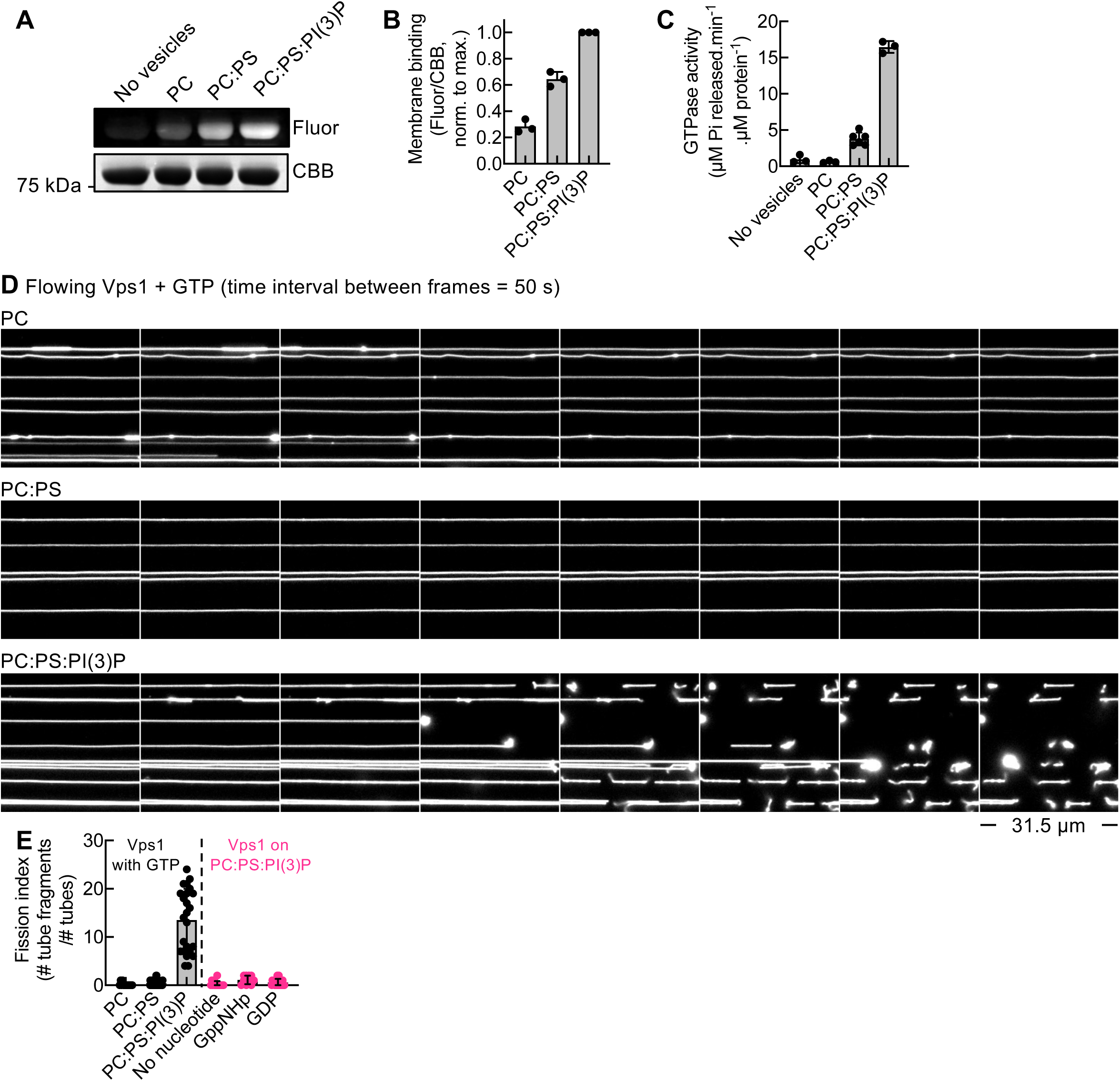
Vps1-catalyzed fission of tubular membrane templates. (A) Representative gel from a PLiMAP experiment showing fluorescence (Fluor) and Coomassie Brilliant Blue (CBB) staining of Vps1 in the absence or presence of vesicles of the indicated composition. (B) Fluor/CBB ratio normalized to the signal seen with PC:PS:PI(3)P (maximum). Data represent the mean ± SD of 3 experiments. (C) GTPase activity of Vps1 in the absence or presence of vesicles of the indicated composition. Data represents the mean ± SD of at least 3 experiments. (D) Representative images from a time-lapse movie acquired after flowing Vps1 with GTP onto SMrTs of the indicated composition. (E) Fission index on templates of the indicated lipid composition and with different nucleotides. Data represents mean ± SD of at least 10 microscope fields across 2 experiments.

### Mechanism of Vps1-catalyzed membrane fission

Fission dynamins form curved helical scaffolds around tubular membranes and their ability to do so is therefore expected to be influenced by the membrane curvature or topology of the tube (Kamerkar *et al*., 2018; Ford and Chappie, 2019; Jimah and Hinshaw, 2019; Khurana *et al*., 2023; Sarkar and Pucadyil, 2025). This has relevance to physiology since membrane tubulating proteins like the SNX-BAR proteins involved in the retromer pathway sculpt membranes to form tubules of a defined curvature (Kovtun *et al*., 2018). Since SMrTs display tubes of varying sizes, we assessed the range of tube that can be severed by Vps1. Imaging a representative field of SMrTs before and after flowing Vps1 with GTP showed that tubes of ∼20 nm radius were severed but those above 40 nm radius remained intact (Fig. 2A). To quantify this observation further, we scored the fraction of tubes showing at least one cut (fission probability) and plotted it against the starting tube size. Such a plot reveals that tubes below 30 nm radius showed a high fission probability, which dropped precipitously on tubes of wider radii (Fig. 2B). To better understand this process, we analyzed a movie of a thin (∼15 nm radius) and a thick (∼40 nm radius) tube as they were exposed to Vps1 with GTP (Movie S2). Time-lapse images from this movie show localized dimming of tube fluorescence at early time points upon flowing Vps1 with GTP (Fig. 2C, white arrowheads). Since the tubes are diffraction-limited objects, dimming of fluorescence implies thinning or constriction of the tube. These constrictions appeared on both the thin and the thick tube, likely signifying attempts made by the Vps1 scaffold to sever these tubes. Thus, it’s reasonable to assume that the Vps1 scaffold functions similarly on tubes within a size range of 15-40 nm radius. But only the thin tube gets severed (Fig. 2C, Movie S2). The constricted intermediates generated on the tube were transient, which precluded further analysis. We therefore resorted to reconstituting the Vps1-catalyzed fission reaction in a stage-wise manner by first flowing Vps1 with GppNHp, which would reflect the GTP-bound state and then flowing GTP to exchange the bound GppNHp for GTP.

**Fig. 2.**
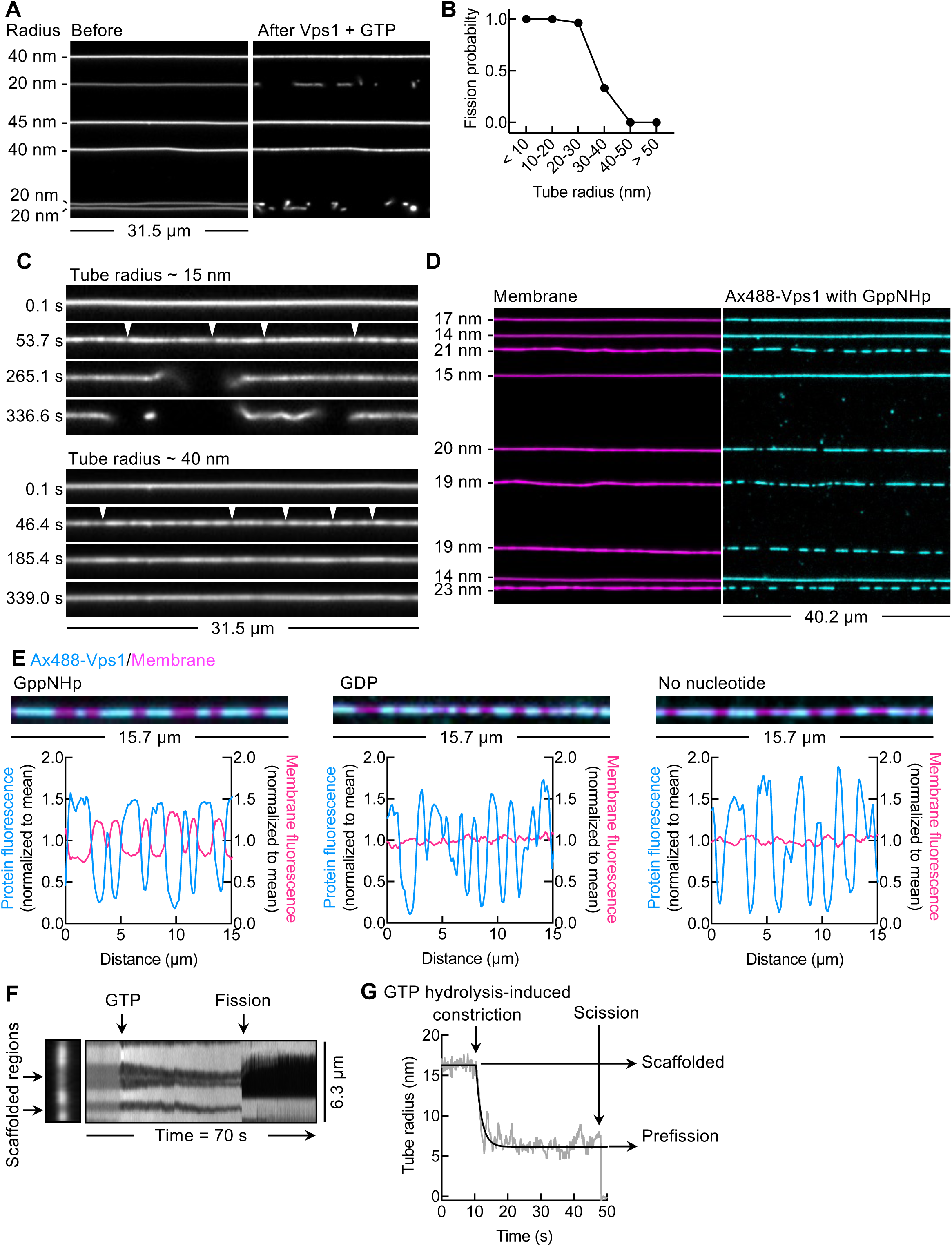
Mechanism of Vps1-catalyzed membrane fission. (A) Representative images of SMrTs before and after flowing Vps1 with GTP. (B) Fission probability, defined as the fraction of tubes showing at least one cut, as a function of tube radius. Data represent probabilities estimated for 5 tubes of <10 nm, 70 tubes of 10-20 nm, 28 tubes of 20-30 nm, 3 tubes of 30-40 nm, 2 tubes of 40-50 nm and 3 of > 50 nm radius. (C) Images from a time-lapse movie acquired after flowing Vps1 with GTP on a thin (∼15 nm radius) and thick (∼40 nm radius) tube. White arrowheads mark localized sites of constriction. (D) Representative image of SMrTs showing Ax488-labeled Vps1 on tubes in the presence of GppNHp. (E) Representative images and associated line profiles of a single tube showing Ax488-labeled Vps1 with GppNHp, GDP and in the absence of nucleotides. (F) Kymograph from a movie of a single tube with GppNHp-bound Vps1 scaffolds responding to GTP addition. (G) Plot showing the tube radius under a GppNHp-bound Vps1 scaffold responding to GTP addition. Data are fitted to a plateau followed by a one- phase exponential decay.

Fluorescently labeled Vps1 (Ax488-Vps1) mixed with GppNHp were uniformly distributed on some tubes but appeared as discrete assemblies, likely reflecting scaffolds, on others (Fig. 2D). This heterogeneity arises from the starting tube size because the distribution appeared more discrete on tubes above ∼15 nm radius (Fig. 2D). The scaffolded regions coincided with lower tube fluorescence than seen on adjacent regions of the same tube lacking scaffolds (Fig. 2E and associated line profiles), indicating constriction of the underlying tube. Similar scaffolds were also seen with GDP and in the absence of nucleotides, but they did not display constriction (Fig. 2E and associated line profiles). These results indicate that GppNHp-bound Vps1 scaffolds are membrane active and constrict the underlying tube. The estimated tube radius under such scaffolds was 19.3 ± 2.1 nm (mean ± S.D., n = 43 scaffolds). Flowing GTP on to GppNHp-bound Vps1 scaffolds caused a sudden further constriction of the underlying tube, which is evident in a kymograph acquired in the membrane fluorescence channel (Fig. 2F, Movie S3). Remarkably, the highly constricted intermediate persisted for a long duration before fission (Fig. 2F, Movie S3), which is quite unlike that seen with the mammalian endocytic and mitochondrial dynamins where the GTPase-induced constriction leads to immediate fission (Dar and Pucadyil, 2017; Pemberton *et al*., 2025). Tracing the size of the scaffolded tube shows that GTP hydrolysis causes constriction to an intermediate of ∼7 nm radius, prior to scission (Fig. 2F). The estimated tube radius of such a prefission intermediate from several independent fission events was 7.1 ± 1.8 nm (mean ± S.D., n = 20 events).

Scaffolds are formed by the tendency of Vps1 to self-assemble and the resultant stimulation in GTPase activity is required to further constrict the tube for fission. These results explain why self-assembly and stimulated GTPase activities are both necessary requirements for fission. Based on the observed pathway to fission, we reason that the inability for a thick tube to undergo fission lies in the inability for the Vps1 scaffold-induced constriction to attain the critical prefission intermediate on the thick tube. Together, these results delineate how Vps1 catalyzes fission and explain why fission displays a substrate-size dependence.

### Vps1-catalyzed fission is required for Vps10 trafficking

Vps10 tranics between the endosome or PVC and the TGN and its retrieval from the PVC requires the retromer complex and Vps1. Loss of Vps1 causes the mis-sorting of Vps10 to the vacuole and represents a functional read-out of the retromer trafficking pathway (Marcusson *et al*., 1994; Nothwehr *et al*., 1995; Cooper and Stevens, 1996; Arlt *et al*., 2014; Chi *et al*., 2014). We generated strains with Vps10 fused with mNeonGreen (Vps10-mNG) and the vacuolar marker, V-ATPase subunit Vph1 fused with mScarlet (Vph1-mSc) at their endogenous loci. Widefield imaging of wildtype cells showed Vph1-mSc marking large vacuoles that were devoid of Vps10-mNG (Fig. 3A). Vps10-mNG was found as foci (Fig. 3A, Movie S4), likely reflecting its localization to the TGN and endosomes (Chi *et al*., 2014). In contrast, *vps1Δ* cells showed multi-lobed vacuoles (Fig. 3B), which has been referred to as the class F vacuolar phenotype (Raymond *et al*., 1992; Rooij *et al*., 2010). *vps1Δ* cells displayed Vps10-mNG on vacuoles indicating mis-sorting (Fig. 3B, Movie S5). We introduced a *VPS1\** transgene in the *vps1Δ* background to test Vps1 functions through a rescue strategy. This transgene has VPS1 fused to the same tags as those used for in vitro studies in addition to a 13 residue-long ALFA tag at the C-terminus (see below). In *vps1Δ* cells expressing *VPS1\**, vacuole morphology was largely like that seen in wildtype and Vps10-mNG signals appeared punctate and distinct from the vacuole (Fig. 3C, Movie S6), implying rescue of both the vacuole morphology and Vps10-mNG mis-sorting. To analyze Vps10 mis-sorting at the population level, we used an image analysis routine that estimates the Vps10-mNG and Vph1-mSc fluorescence within a mask generated using the Vph1-mSc signal. Ratio of Vps10-mNG to Vph1-mSc fluorescence within the mask reports on the vacuolar Vps10-mNG. Such analyses revealed higher levels of Vps10-mNG on vacuoles in *vps1Δ* compared to WT, which was rescued in *VPS1\** (Fig. 3D). These results establish a quantitative vacuolar localization assay to monitor Vps1 functions. Furthermore, the *vps1Δ* strain shows growth defects at high temperature, which was rescued with *VPS1\** (Fig. S1) (Rothman *et al*., 1990).

**Fig. 3.**
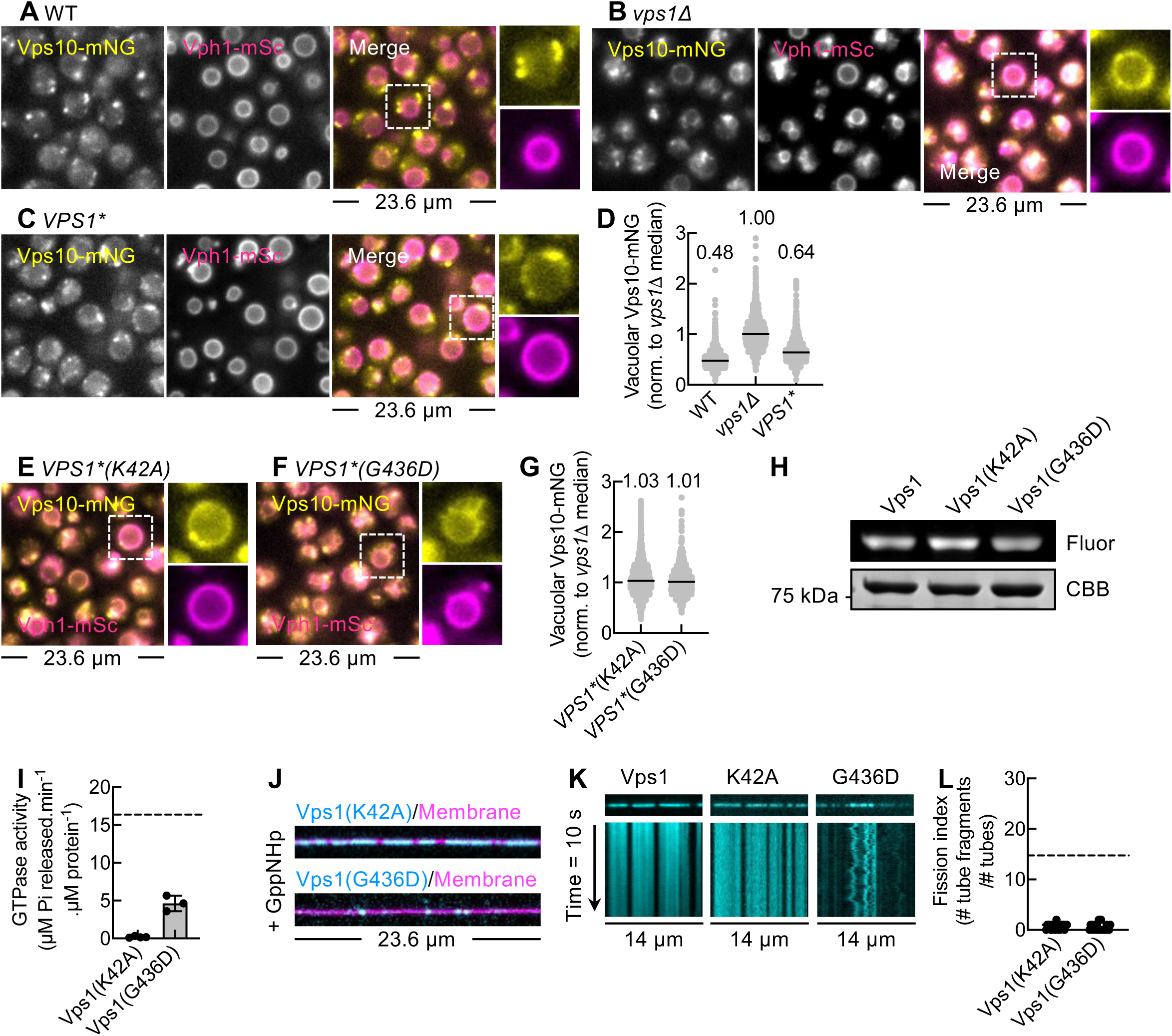
Vps1 functions in Vps10 traWicking. Representative fluorescence images showing Vps10-mNG and Vph1-mSc in wildtype (WT) (A), *vps1Δ* (B) and *VPS1\** (C). (D) Quantitation of vacuolar Vps10-mNG in 1502 Vph1-mSc-positive particles for WT, 1146 for *vps1Δ* and 1381 for *VPS1\** from 2 experiments. Representative fluorescence images showing Vps10-mNG and Vph1-mSc in *VPS1\**(K42A) (E) and *VPS1\**(G436D) (F) cells. (G) Quantitation of vacuolar Vps10-mNG in 1699 Vph1-mSc-positive particles for *VPS1*(K42A)* and 1021 for *VPS1*(G436D)* from 2 experiments. (H) Representative gel from a PLiMAP experiment showing fluorescence (Fluor) and Coomassie Brilliant Blue (CBB) staining of Vps1 mutants on PC:PS:PI(3)P vesicles. (I) GTPase activity of Vps1 mutants in the presence of PC:PS:PI(3)P vesicles. Data represents the mean ± SD of 3 experiments. Dotted line represents the average GTPase activity seen with Vps1 on PC:PS:PI(3)P vesicles. (J) Representative images of Ax488-labeled Vps1(K42A) and Vps1(G436D) on tubes in the presence of GppNHp. (K) Kymographs from movies imaging fluorescent Vps1, Vps1(K42A) and Vps1(G436D) on tubes. (L) Fission index with Vps1(K42A) and Vps1(G436D) with GTP. Dotted line represents the average fission index seen for Vps1. Data represents mean ± SD of at least 20 microscope fields across 2 experiments. For plots showing quantitation of vacuolar Vps10-mNG (D,G), data are normalized to the median Vps10-mNG fluorescence in *vps1Δ* cells. The numbers in the plot represent the median values and black lines represent the median position. Vacuolar levels of Vps10-mNG in VPS1* mutants and VPS1* were significantly dinerent (p<0.0001, unpaired Mann-Whitney test).

We tested previously reported mutants of Vps1 in the Vps10 endosomal sorting assay (Varlakhanova *et al*., 2018). Cells with VPS1* bearing the P-loop mutation K42A in the G domain or the stalk mutation G436D showed distorted vacuoles and an enrichment of Vps10-mNG in vacuoles (Fig. 3E- G), indicating failure to complement the *vps1Δ* defect. To establish correlations between in vivo and in vitro functions of Vps1, we assessed these mutants in cell-free reconstitution assays. PLiMAP assays revealed that both mutants bound PC:PS:PI(3)P vesicles as well as WT (Fig. 3H). As expected, Vps1(K42A) showed an insignificant GTPase activity while Vps1(G436D) showed a GTPase rate of 4.5 min^-1^, which was 3.5-fold lower than seen for WT (Fig. 3I). GppNHp-bound Vps1(K42A) organized as stable scaffolds on membrane tubes (Fig. 3J), like Vps1. This is apparent in kymographs where scaffold positions remained constant over time for both Vps1 and Vps1(K42A) (Fig. 3K). On the other hand, GppNHp-bound Vps1(G436D) appeared as small clusters that moved laterally on the tube (Fig. 3J,K), signifying defects in scaffolding. Unlike PLiMAP assays, visualizing proteins on membrane tubes involves buffer washes. The high binding avidity resulting from multivalent interactions between oligomerized Vps1 in the scaffold and membrane lipids render it stable to buffer washes. Self-assembly defects would hinder the formation of multivalent interactions and self-assembly mutants would display lower binding avidity, thereby making them prone to dissociation from the membrane upon buffer washes. This would result in low protein densities on the tubes, like seen here for the Vps1(G436D) mutant. Both mutants were defective in fission thus underscoring the importance of self-assembly and stimulated GTPase activities (Fig. 3L). Together, these results correlate Vps10 mis-sorting with defects in Vps1-catalyzed membrane fission.

### Distribution and dynamics of Vps1 in cells

The short ALFA tag when co-expressed with the ALFA nanobody fused with mNeonGreen (ALFA- NB-mNG) allows visualizing Vps1* using the ALIBY strategy, which has been successfully applied to image proteins that function as oligomers (Götzke *et al*., 2019; Akhuli *et al*., 2022). This strategy was used because direct C-terminal fusion to GFP has been reported to perturb Vps1 functions (Suzuki *et al*., 2021; Arlt *et al*., 2023). Furthermore, the growth defect in *vps1Δ* strain at high temperature was rescued with *VPS1\** expressing the ALFA-NB-mNG indicating that binding to ALFA-NB-mNG does not compromise Vps1 functions (Fig. S1).

In cells expressing Vps1*, the ALFA-NB-mNG signal appeared as dynamic foci, some of which were found on the vacuole (Movie S7). On the other hand, Vps1*(K42A) appeared as large and stable foci (Movie S8) while Vps1*(G436D) did not form foci and appeared diffuse in the cytoplasm (Movie S9). We quantitated these observations through a Vps1 foci formation assay by analyzing the coefficient-of-variation (COV) of ALFA-NB-mNG fluorescence (Fig. S2). Vps1* expressing cells showed a COV that was higher than the control not expressing Vps1*. Vps1*(K42A) expressing cells displayed an even higher COV while those expressing Vps1*(G436D) showed a lower COV than seen for Vps1* (Fig. S2). This result correlates well with the observation that Vps1*(K42A) forms larger foci while Vps1*(G436D) does not form foci. Western blots using the ALFA antibody confirmed similar levels of expression of Vps1* and mutants (Fig. S3). The large and stable Vps1*(K42A) foci likely represent self-assemblies trapped in the GTP-bound state while the diffuse Vps1*G436D fluorescence reflects an inability to self-assemble. The small and dynamic Vps1* foci therefore reflects a steady state between self-assembly and GTPase-induced disassembly and dissociation. Thus, the intrinsic ability to self-assemble into scaffolds and display stimulated GTPase activity contributes to organizing Vps1 as foci for optimal function.

### Functional analysis of Insert B in Vps1

Endocytic dynamins engage with phosphoinositide lipids through the PHD, which is present as the membrane-binding foot (Khurana and Pucadyil, 2023). Several dynamins lack a PHD and instead have regions like the Paddle Domain (PD) in BDLP and Opa1, Variable Domain (VD) in Drp1, L4 loop in MxA, and Insert B (InsB) in Vps1 (Jimah and Hinshaw, 2019). InsB has been reported to bind lipids (Smaczynska-de Rooij *et al*., 2019), but a comprehensive understanding of how this region contributes to Vps1 functions is lacking. We therefore analyzed Vps1 mutated in specific regions of InsB using the suite of in vitro and in vivo assays described above.

In Vps1, InsB spans residues 536-612 and sequence alignment of this region across fungi reveals several conserved stretches (Fig. 4A) (Smaczynska-de Rooij *et al*., 2019). The region towards the C-terminus displays a phenylalanine-rich 4F motif (579-587) and a lysine-rich 3K motif (591-593). Since both aromatic and positively charged residues are frequently found in membrane binding regions in proteins, we generated constructs deleted in the 4F motif (Δ4F) and the 3K motif (Δ3K) (Fig. 4A). We also deleted the entire InsB region and replaced it with a short linker (ΔInsB). PLiMAP assays showed that Δ4F bound PC:PS:PI(3)P vesicles as well as full-length Vps1 while Δ3K showed a 40% reduction in binding (Fig. 4B,C), indicating that the 3K motif is directly involved in membrane binding. Deletion of the 3K motif reduced binding to levels like that seen for Vps1 on PC:PS vesicles (Fig. 4C, red dotted line), indicating that the 3K motif accounts for PI(3)P binding. ΔInsB only showed a marginal further reduction in binding, indicating that the 3K motif is the sole lipid binding site in InsB. But the binding of ΔInsB was higher than seen for Vps1 on PC vesicles (Fig. 4C, black dotted line), which suggests the presence of additional sites outside of InsB that can bind PS. Consistent with a PI(3)P-binding defect, Δ3K showed no stimulated GTPase activity on PC:PS:PI(3)P vesicles (Fig. 4D). But Δ4F, which bound PC:PS:PI(3)P vesicles as well as full length Vps1 (Fig. 4B,C), was also defective in GTPase activity. This is reminiscent of what was observed with the self-assembly defective G436D mutant (Fig. 3H,I). Indeed, imaging fluorescent Δ4F on membrane tubes revealed substantially lower levels of protein like was seen for Vps1(G436D) (Fig. 4E).

**Fig. 4.**
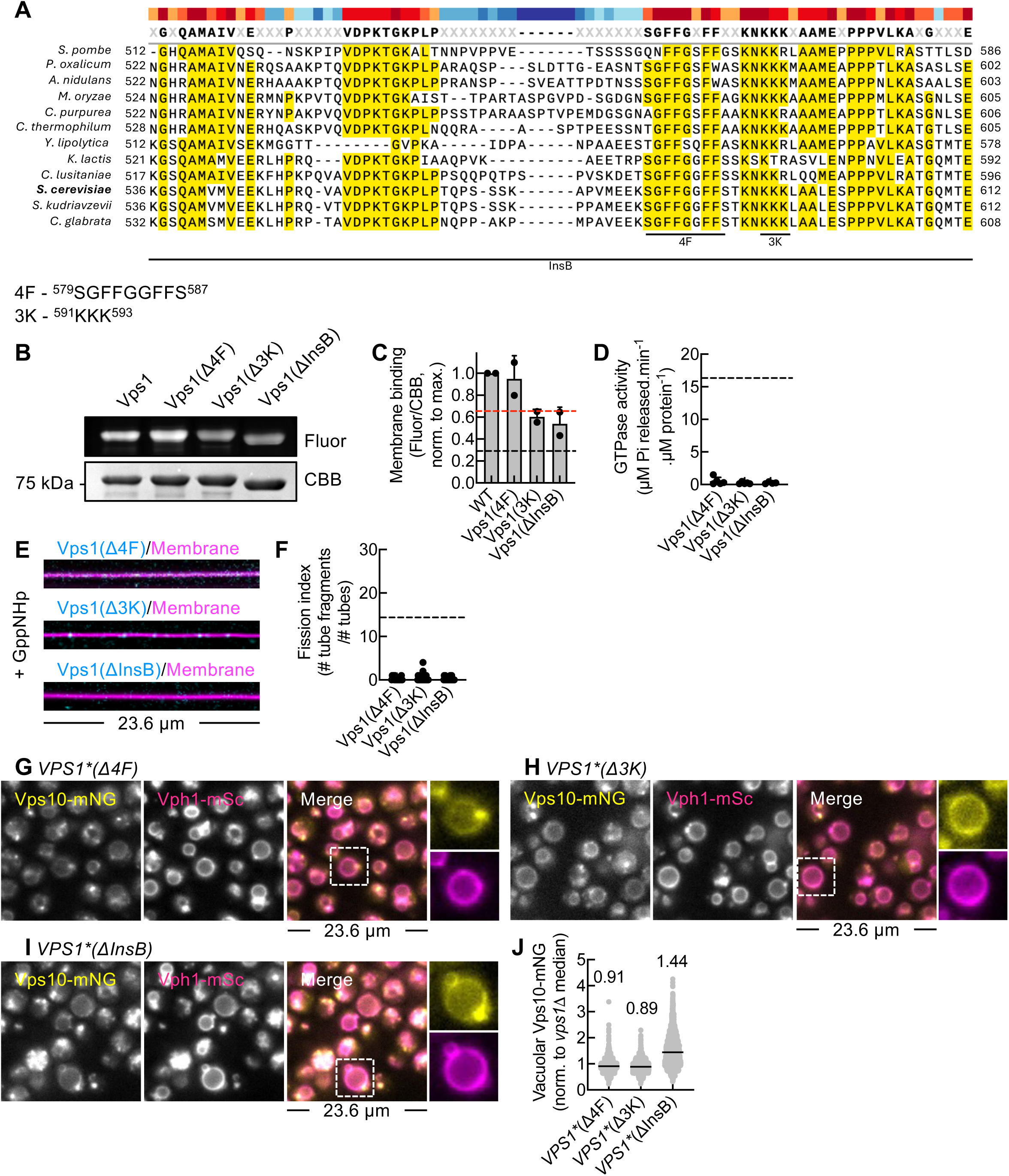
Functions of Insert B in Vps1. (A) Sequence alignment of InsB from various fungi. Amino acid sequences of the 4F and 3K motifs in *S. cerevisiae* are shown. (B) Representative gel from a PLiMAP experiment showing fluorescence (Fluor) and Coomassie Brilliant Blue (CBB) staining of InsB mutants in presence of PC:PS:PI(3)P vesicles. (C) Fluor/CBB ratio of InsB normalized to the signal seen for WT. Data represent the mean ± SD of 2 experiments. Red and black dotted lines mark the average Fluor/CBB ratio seen for WT on PC:PS and PC vesicles, respectively. (D) GTPase activity of InsB mutants on PC:PS:PI(3)P vesicles. Data represent the mean ± SD of at least 5 experiments. Black dotted line marks the average GTPase activity of Vps1. (E) Representative images of Ax488-labeled InsB mutants on tubes in the presence of GppNHp. (F) Fission index with InsB mutants in the presence of GTP. Data represents mean ± SD of at least 13 microscope fields across 2 experiments. Dotted line marks the average fission index seen for Vps1. Representative fluorescence images showing Vps10-mNG and Vph1-mSc in *VPS1*(Δ4F)* (G), *VPS1*(Δ3K)* (H) and *VPS1*(ΔInsB)* (I). (J) Quantitation of vacuolar Vps10-mNG fluorescence in 1844 Vph1-mSc-positive particles for *VPS1*(Δ4F)*, 1624 for *VPS1*(Δ3K)* and 2928 for *VPS1*(ΔInsB)* from 2 experiments. Data are normalized to the median Vps10-mNG fluorescence in *vps1Δ* cells. The numbers in the plot represent the median values and black lines represent the median position. Vacuolar levels of Vps10-mNG in VPS1* mutants and VPS1* were significantly dinerent (p<0.0001, unpaired Mann-Whitney test).

As expected, Δ3K and ΔInsB, where both the 4F and 3K motifs are absent, also showed no binding to tubes (Fig. 4E). Because of these defects, Δ4F, Δ3K and ΔInsB lacked fission activity (Fig. 4F).

Next, we analyzed these mutants for their distribution and functions in Vps10 trafficking.

Western blots showed similar levels of expression of these constructs (Fig. S3). Imaging ALFA-NB-mNG revealed a heterogenous distribution of Vps1*(Δ4F) and Vps1*(Δ3K) in cells. In some cells, they were organized as small foci while in others, no foci were apparent (Movies S10,S11), indicating self-assembly defects. This was more apparent with Vps1*(Δ4F) than with Vps1*(Δ3K). On the other hand, Vps1*(ΔInsB) was diffusely distributed (Movie S12), like Vps1*(G436D), indicating that the combined loss of the 4F and 3K motifs results in a severe self-assembly defect. These interpretations were corroborated through the Vps1 foci formation assay where Vps1*(Δ4F) and Vps1*(Δ3K) showed a lower COV than wildtype but higher COV than controls while Vps1*(ΔInsB) showed a COV that was like that seen for Vps1*(G436D) (Fig. S2). Finally, cells expressing the Δ4F, Δ3K and ΔInsB mutants were unable to rescue Vps10-mNG mis-localization, which is apparent in both the representative images (Fig. 4G-I) and from the quantitative vacuolar localization assay (Fig. 4I). Together, these results parse out independent attributes among the conserved motifs in InsB. The 3K motif mediates PI(3)P binding while the 4F motif facilitates self-assembly and both these motifs are necessary for Vps1 functions.

### Quantitative vacuolar proteomics reveals insights into Vps1-dependent cargo trafficking and organelle quality control pathways

Our results correlate Vps1 functions in membrane fission to the fidelity of trafficking of the retromer cargo Vps10. Since membrane fission is a necessary step for the formation of transport carriers, Vps1 functions are likely required for the trafficking of other cargo. Based on the logic that trafficking defects result in vacuolar enrichment of Vps10, we analyzed proteome-wide alterations in vacuoles from *vps1Δ* and wildtype cells.

Vacuoles were isolated on a density gradient and processed for quantitative proteomics. Filtering data based on the number of peptides detected and the confidence level of peptide identification gave us a list of 284 total proteins (Table S1). Vacuolar proteins were the most enriched among organellar proteins in this list and validates the vacuole isolation procedure (Table S2). Using a screening criterion of ≥ 2-fold change in abundance, we find that a significant fraction (∼67%) of the detected proteins were depleted in the *vps1Δ* vacuole while only a small (∼4%) of the proteins were enriched (Table S1). Vps1 is therefore critical for maintaining the vacuolar proteome. Soluble vacuole-resident enzymes that are transported by Vps10 like Ecm14, Pep4, Npc2, Ape3, Atg42 and Prc1 were depleted while Vps10 itself was enriched in the *vps1Δ* vacuole (Fig. 5A, Table S1) (Eising *et al*., 2022). These results corroborate the model for Vps10-dependent protein sorting into vacuoles (Marcusson *et al*., 1994; Nothwehr *et al*., 1995; Cooper and Stevens, 1996; Arlt *et al*., 2014; Chi *et al*., 2014). Gene ontology analysis revealed that several proteins involved in vacuole fusion, amino acid metabolism and vacuole organization were significantly depleted in *vps1Δ* vacuoles (Table S3). This included the Rab Ypt7, which is required for vacuole fusion and retromer-dependent trafficking (Hickey *et al*., 2009; Balderhaar *et al*., 2010; Liu *et al*., 2012). Other Rabs like Vps21, Ypt1 and Ypt32 were also depleted in *vps1Δ* vacuoles. The stress response kinase Tor1 was also depleted in *vps1Δ* vacuoles (Table S1). Ape1, which utilizes the Cytoplasm-to-Vacuole Targeting (Cvt) pathway to reach the vacuole, was also depleted in *vps1Δ* vacuoles (Table S1), indicating that Vps1 might regulate flux in the CVT pathway. Several integral membrane proteins were depleted in *vps1Δ* vacuoles. These include proteins like Fcy2, Dip5, Yvc1 and Fui1 that utilize the GGA pathway and proteins like Zrt3, Vtc2, Vtc3, Vtc4, Pho91, Syg1, Pho8, Ybt1, Ypk9, Vnx1, Pff1, Smf3, Fmp42 and Ccc1 that utilize the AP-3 pathway for trafficking from the TGN to the vacuole (Table S1) (Eising *et al*., 2022). To confirm results from the proteomic analysis and understand potential mechanisms, we analyzed the phosphatidylcholine transporter Ybt1 and the zinc transporter Zrc1. We generated strains expressing Ybt1-mNG or Zrc1-mNG with Vph1-mSc. Both Ybt1-mNG (Fig. 5B) and Zrc1-mNG (Fig. 5D) co-localized well with Vph1-mSc in wildtype and *vps1Δ* cells, indicating no apparent sorting defect due to the absence of Vps1. Consistent with the proteomic analysis, quantitative vacuolar localization analysis showed significant depletion of Ybt1-mNG (Fig. 5C) and Zrc1-mNG (Fig. 5E) levels in wildtype and *vps1Δ* cells.

**Fig. 5.**
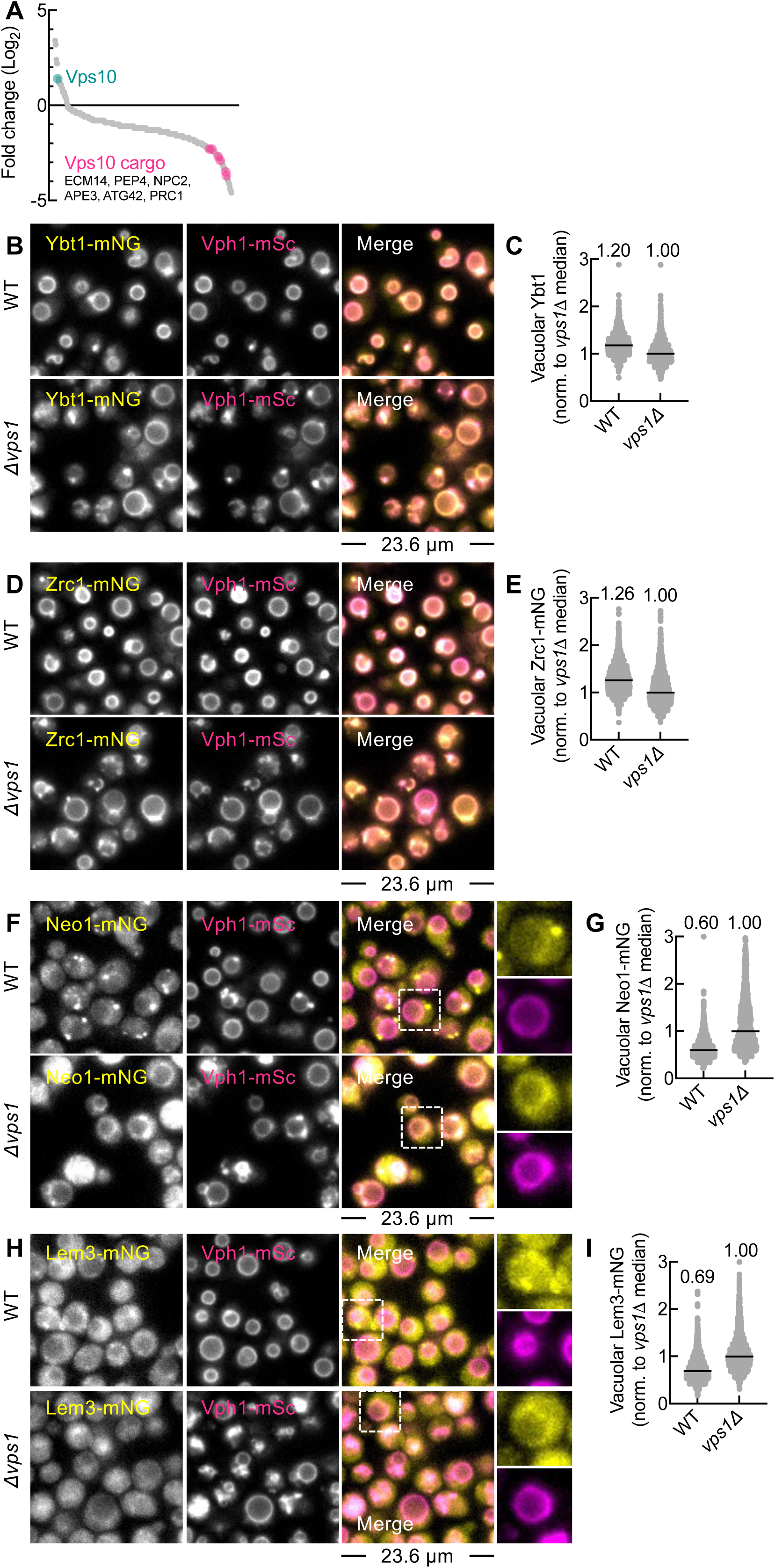
Quantitative proteomics of *vps1Δ* vacuoles. (A) Plot showing the 284 proteins (grey) detected through quantitative proteomics and sorted based on their abundance in *vps1Δ* vacuoles. Vps10 (green) and Vps10 cargo (red) are marked. (B) Representative fluorescence images showing Ybt1-mNG against Vph1-mSc in wildtype and *vps1Δ* cells. (C) Quantitation of vacuolar Ybt1-mNG fluorescence in 1333 particles for WT and 1465 for *vps1Δ* from 2 experiments. (D) Representative fluorescence images showing Zrc1-mNG against Vph1-mSc in wildtype and *vps1Δ* cells. (E) Quantitation of vacuolar Zrc1-mNG fluorescence in 2421 particles for WT and 2473 for *vps1Δ* from 2 experiments. (F) Representative fluorescence images showing Neo1-mNG and Vph1-mSc in WT and *vps1Δ* cells. (G) Quantitation of vacuolar Neo1-mNG fluorescence in 2110 Vph1-mSc-positive particles in WT and 2179 for *vps1Δ* cells from 2 experiments. (H) Representative fluorescence images showing Lem3-mNG and Vph1-mSc in WT and *vps1Δ* cells. (I) Quantitation of vacuolar Lem3-mNG fluorescence in 2101 Vph1-mSc-positive particles in WT and 2774 in *vps1Δ* cells from 2 experiments. For plots showing quantitation of vacuolar cargo (C,E,G,H), data are normalized to the median cargo fluorescence in *vps1Δ* cells. The numbers in the plot represent the median values and black lines represent the median position. Levels of all cargo between WT and *vps1Δ* cells were significantly dinerent (p<0.0001, unpaired Mann-Whitney test).

In addition to Vps10, *vps1Δ* vacuoles were enriched in the Golgi-localized V-ATPase subunit Stv1 and the Golgi-localized phospholipid flippases Neo1 and Dnf2, and its adaptor Lem3 (Table S1). Unlike the vacuole localized Vph1, Stv1 traffics between the TGN and endosomes and is mis-sorted to the vacuole upon loss of retromer functions, indicating that Stv1 is a retromer cargo (Finnigan *et al*., 2011). Neo1 and its mammalian homolog ATP9A have recently been reported as retromer cargo (Dalton *et al*., 2017; McGough *et al*., 2018; Suzuki *et al*., 2021). Microscopic analysis revealed Neo1-mNG as foci that were distinct from the vacuole in wildtype cells (Fig. 5F, Movie 13). These foci were absent in *vps1Δ* cells and Neo1-mNG appeared localized to the vacuole (Fig. 5F, Movie 14). Indeed, quantitative vacuolar localization analysis verified enrichment (Fig. 5G). Proteomic analysis also revealed Lem3 and Dnf2 to be enriched in *vps1Δ* vacuoles. Recent work shows that Dnf1/2 are trafficked by the retromer from the endosome or PVC to the TGN (Jiménez *et al*., 2024). Dnf1/2 are found in a complex with Lem3 (Bai *et al*., 2020). Furthermore, Dnf1/2 and Lem3 have been reported to interact with the Golgi-associated retrograde protein (GARP) complex, which functions in the retrograde transport of proteins from the PVC or endosome and vacuolar levels of Dnf1/2 and Lem3 show an increase upon depletion of the GARP complex (Conibear *et al*., 2003; Eising *et al*., 2019). Lem3-mNG appeared as faint foci in wildtype cells but was more diffuse in *vps1Δ* cells (Fig. 5H, Movie S15, S16). Quantitative vacuolar localization analysis showed a significant enrichment in *vps1Δ* vacuoles (Fig. 5I). *vps1Δ* vacuoles were also enriched in the NADH-cytochrome b5 reductase Cbr1 (Table S1). Cbr1 has an N-terminal transmembrane anchor and is involved in sterol biosynthesis and lipid peroxidation (Hall *et al*., 2022). Thus, loss of Vps1 enriches the vacuole with a small set of proteins that represent cargo for the retromer pathway while depleting the vacuole of the bulk of proteins.

## Discussion

Using a combination of biochemistry, cell-free reconstitution and live cell imaging, we comprehensively analyze functions of the yeast dynamin Vps1. We find Vps1 to be sufficient for constricting tubular membranes, thereby causing fission and that this activity correlates well with endosomal protein sorting. In mammals, the endocytic dynamin has been proposed to function along with the WASH complex to facilitate fission of retromer-containing tubular intermediates generated on endosomes (Derivery *et al*., 2009). Thus, the involvement of dynamins in endosomal protein sorting and trafficking appears to be conserved. Vps1 is linked to other pathways that involve membrane remodeling, such as endocytosis, autophagy and peroxisome division (Kuravi *et al*., 2006; Rooij *et al*., 2010; Palmer *et al*., 2015; Arlt *et al*., 2023). By extension, it is quite likely that Vps1 functions similarly in catalyzing fission in these pathways.

From the membrane side, fission appears to be regulated by membrane topology while from the protein side, fission requires Vps1 to self-assemble into scaffolds, which in turn stimulates its GTPase activity thereby forcing membrane constriction to cause fission. Our results indicate that PI(3)P stabilizes Vps1 on the membrane. Previous structural and biochemical analyses on lipid nanotubes revealed a propensity for Vps1 to self-assemble into scaffolds but surprisingly no significant stimulation in GTPase activity was observed (Varlakhanova *et al*., 2018). The reported GTPase rates are in fact like what we find here with Vps1 on PC:PS vesicles. Our results instead reveal a ∼30-fold stimulation in GTPase activity.

We are unsure of the cause for this difference, but it may reflect the intrinsic properties of the *C. thermophilum* Vps1 tested earlier compared to the *S. cerevisiae* tested here. Vps1 binds PI(3)P through a conserved lysine-rich (3K) motif located in InsB and self-assembles into scaffolds around tubes, determinants for which lie within the stalk but surprisingly are also encoded in a conserved phenylalanine-rich (4F) motif again present in InsB. GTP-bound Vps1 scaffolds are by themselves membrane active and constrict tubes to a radius of ∼20 nm. Cryo-EM reconstructions of GTP-bound Vps1 scaffolds formed in solution reveal a lumen radius of ∼10 nm (Varlakhanova et al., 2018), which after accounting for the bilayer thickness is significantly smaller than the ∼20 nm tube radius seen here. This indicates that the membrane acts as a barrier to restrict the intrinsic curvature of Vps1 scaffolds. GTP hydrolysis causes further constriction of the Vps1 scaffold, leading to thinning of the underlying tube, which upon reaching a critical ∼7 nm radius, undergoes fission. Membrane topology regulates Vps1 functions by restricting its fission activity to tubes below ∼30 nm radius. Based on our previous analysis, a fission probability cut-off of 30 nm tube radius places Vps1 functions closer to the mammalian endocytic dynamin than the mitochondrial dynamin (Kamerkar *et al*., 2018; Khurana *et al*., 2023; Sarkar and Pucadyil, 2025). This substrate size cut-off arises not from the inability of Vps1 to organize into scaffolds but rather from a limitation in reaching the critical radius on thick tubes. Vps1 functions in concert with the membrane tubulating SNX-BAR proteins on endosomes. SNX-BAR-coated membrane tubules display a radius of ∼15 nm (Kovtun *et al*., 2018), which falls within the size range for Vps1 to catalyze fission.

Our results indicate that InsB harbors determinants both for membrane binding and self-assembly. Previous results from sedimentation assays showed that mutating the 3K motif affects binding to PI(4,5)P2 but not PI(3)P lipids and that the 3K motif was not important for Vps1 functions in endosomal protein sorting (Smaczynska-de Rooij *et al*., 2019). Using a sensitive membrane-binding PLiMAP assay, our results indicate otherwise. The 3K motif coordinates binding to PI(3)P lipids and is likely the sole membrane binding motif in InsB because both the Δ3K and the ΔInsB mutants show similar levels of reduction in binding to PC:PS:PI(3)P membranes. A similar lysine-rich motif is also found in the mammalian mitochondrial Drp1, which is involved in binding to cardiolipin (Bustillo-Zabalbeitia *et al*., 2014). We find residual membrane binding with the ΔInsB mutant, suggesting membrane interactions outside of the InsB. We also identify a novel 4F in InsB that functions in self-assembly. Deletion of this motif showed severe loss of Vps1 functions in vitro, in a manner that is consistent with a self-assembly defect. However, both mutants (Δ4F and Δ3K) were found to organize as smaller foci in some cells, which could reflect their localized recruitment via adaptor-based interactions, which are not recapitulated in the in vitro assays. Importantly, these mutants are unable to function in endosomal protein sorting thus emphasizing their importance to Vps1 functions.

Quantitative vacuolar/lysosome proteomics has been previously used to analyze trafficking pathways (Eising *et al*., 2019, 2022; Daly *et al*., 2023). But to the best of our knowledge, such an approach to evaluate protein trafficking upon loss of a membrane fission protein has not been reported. Our analysis reveals novel insights into proteome-wide alterations that accrue upon loss of Vps1. Disruption of retromer functions have been reported to cause an enrichment of Rab7 levels on mammalian lysosomes (Daly *et al*., 2023). In contrast, our results indicate a reduction in Ypt7 levels on *Δvps1* vacuoles. We are unsure of the mechanism as the proteomics data do not identify GAPs or GEFs of Ypt7. Ypt7 is a major regulator of fusion and retromer-dependent trafficking on the vacuole (Hickey *et al*., 2009; Balderhaar *et al*., 2010; Liu *et al*., 2012). Its depletion upon loss of Vps1 functions could signify a feedback mechanism wherein fission in turn regulates upstream reactions involving coat protein assembly. Loss of Vps1 causes depletion of Tor1, which is consistent with results seen in mammalian cells upon loss of retromer functions (Daly *et al*., 2023). Since Tor1 regulates stress response pathways, its depletion could explain the growth defect seen in *vps1Δ* cells at high temperatures. Furthermore, loss of Vps1 results in a general deficiency in the levels of vacuolar proteins. The AP-3 pathway delivers the bulk of proteins for vacuole biogenesis (Eising *et al*., 2022) and the loss of AP-3 causes the rerouting of vacuolar proteins to the plasma membrane via the secretory pathway, like that seen upon loss of Vps1 (Nothwehr *et al*., 1995; Dell’Angelica *et al*., 1999). Interestingly, the combined deletion of AP-3 and Vps1 causes synthetic lethality (Stepp *et al*., 1997), which suggests that trafficking pathways that become reconfigured upon loss of AP-3 or Vps1 must converge at some point. In addition, or alternatively, the reduced levels of vacuolar proteins could arise from defects in fusion due to the loss of Ypt7 or enhanced instability and degradation. Importantly, loss of Vps1 results in the enrichment of Vps10, Neo1, Dnf2, Lem3 and Stv1. This finding corroborates previous results that have identified these proteins as retromer cargo. While the recycling itinerary of Vps10 manages to deliver newly synthesized proteins to the vacuole, a similar itinerary seen for the phospholipid flippases Neo1 and Dnf1/2 signifies the important role played by the retromer pathway in regulating phospholipid asymmetry in the endosome and TGN (Sebastian *et al*., 2012).

Vps1 appears to be critical in regulating the trafficking of proteins both into and out of the vacuole and its loss results in global alterations in the vacuolar proteome. Vacuolar enrichment of retromer cargo upon loss of Vps1 can be rationalized based on our understanding of the retromer trafficking pathway. However, reasons for depletion of the bulk of vacuolar proteins remain unclear. Genetic deletions cause trafficking pathways to become reconfigured and the collateral effects arising from these alterations accrue over several generations. Considering these caveats, perhaps a better strategy is to analyze protein trafficking upon acute loss of Vps1 functions and is an aspect that awaits further investigation. Vps1 was identified in a genetic screen for factors involved in vacuolar protein sorting. Here, we establish a quantitative assay that can be applied to large populations to assess mis- sorting of endosomal cargo, which has allowed us to analyze Vps1 functions and validate the trafficking itineraries of other retromer cargos. Defects in retromer functions are linked to a variety of metabolic and neurological disorders (Lane *et al*., 2012; Sebastian *et al*., 2012; Small and Petsko, 2015; Vagnozzi and Praticò, 2019; Cullen *et al*., 2024). Our assays using the yeast model could be applied to test the molecular basis of such pathologies as well as screen small molecule modulators of the retromer pathway and represent an exciting avenue for future research.

## Materials and methods

### DNA constructs

The Vps1 gene was amplified from yeast genomic DNA and cloned into a pET15b vector with an N-terminal 6xHis followed by a TEV cleavage site and a C-terminal StrepII tag. Site-directed mutagenesis was carried out using PCR and confirmed by sequencing. Vps1 and mutants were cloned in pYM16 with an AlFA tag at the C-terminus to generate the *VPS1\** strains. Tables S4 and S5 list the constructs and strains used in this study.

### Protein expression and purification

Vps1 was expressed in T7 express (NEB) using IPTG induction. Transformants were grown to an OD of 0.6-0.8 at 37 °C, induced with 0.1 mM IPTG and left overnight at 22 °C. Cultures were pelleted and stored at -40 °C. Frozen pellets were thawed in 20 mM HEPES pH 7.4, 500 mM NaCl with 1% Triton X-100 and 1% PMSF and lysed by sonication. Lysates were spun at 30,000 g for 20 min and the supernatant was loaded onto a 5 ml Strep-trap HP or Streptactin XT column (GE Lifesciences). The column was washed first with 20 mM HEPES pH 7.4, 500 mM NaCl, then with 20 mM HEPES pH 7.4, 500 mM NaCl containing 100 mM EDTA to remove any bound nucleotides and then again with 20 mM HEPES pH 7.4, 500 mM NaCl. Buffer was exchanged with 20 mM HEPES pH 7.4, 150 mM NaCl and the bound protein was eluted either using 2.5 mM desthiobiotin (Sigma) or 5 mM biotin (HiMedia) in 20 mM HEPES pH 7.4, 150 mM NaCl. Proteins were spun at 100,000 g to remove aggregates before use in assays.

### Vesicle preparation

Dioleoyl phosphatidylcholine (DOPC), dioleoyl phosphatidylserine (DOPS) and phosphatidylinositol-3- phosphate (PI(3)P) were from Avanti Polar Lipids. Lipids were aliquoted in the required proportion into a clean glass tube and dried under vacuum. Lipids were hydrated with deionized water to form vesicles and extruded using a 100-nm pore size filter.

### PLiMAP assays

Proximity-based Labelling of Membrane Associated Proteins (PLiMAP) assays were carried out as described earlier (Jose and Pucadyil, 2020; Jose *et al*., 2020). The fluorescent crosslinker lipid TMR Diazirine PE (TDPE) was synthesized from TopFluor TMR PE (Avanti Polar Lipids) and SDA NHS Diazirine (Thermo Scientific) as described earlier (Jose and Pucadyil, 2020; Jose *et al*., 2020). Proteins were incubated with vesicles at a protein:lipid molar ratio of 1:100 in 20 mM HEPES pH 7.4, 150 mM NaCl for 30 min at room temperature in the dark. Samples were exposed to UV (365 nm, UVP crosslinker CL- 1000L) at an intensity of 200 mJ.cm^-2^ for 1 min and resolved through SDS-PAGE. Gels were first imaged using a laser-based Typhoon scanner for TDPE fluorescence and then stained with Coomassie Brilliant Blue and imaged on an iBright 1500 (Invitrogen). Fluorescence images were analyzed and rendered using Fiji.

### GTPase assays

Proteins (1 μM) were mixed with GTP (1 mM, Jena Biosciences) in the absence or presence of vesicles (100 μM) in 20 mM HEPES pH 7.4, 150 mM NaCl, 1 mM MgCl_2_ and incubated at 30 °C. 20 μl aliquots were taken at regular intervals and quenched with 10 μl of 0.5 mM EDTA (pH 8.0). The released inorganic phosphate was assayed with the Malachite Green reagent (Leonard *et al*., 2005).

### Fluorescence imaging

SMrT templates and yeast cells were imaged through a 100x,1.4 NA oil immersion objective on a motorized Olympus IX73 microscope attached to a pE4000 (CoolLED) light source and an Evolve EMCCD (Photometrics) camera. Image acquisition was controlled by μManager.

### Supported Membrane Templates (SMrT) and image analysis

Supported membrane templates (SMrT) were prepared as described earlier (Dar *et al*., 2017). Lipids and Texas Red DHPE (Thermo Fisher scientific) were aliquoted in the required proportion in a glass vial and diluted to 1 mM total lipid concentration. 1-2 μl of this mix was spotted on a glass coverslip covalently conjugated with PEG400 and dried. The coverslip was assembled in an FCS2 flow cell (Bioptechs) and hydrated using 20 mM HEPES pH 7.4 with 150 mM NaCl. Membrane tubes are formed by passing buffer at high flow rates and leaves behind a supported bilayer at the place where the lipid mix was spotted.

Proteins were introduced at low flow rates at a final concentration of 0.5 μM in the presence of 1 mM nucleotides (Jena Biosciences) and 1 mM MgCl_2_ in 20 mM HEPES pH 7.4, 150 mM NaCl. Time lapse imaging was carried out with buffer containing an oxygen scavenger cocktail (Dar *et al*., 2017; Roy and Pucadyil, 2022). To visualize fluorescently labelled proteins on membrane tubes, 0.5 μM of Vps1 was premixed with 10-fold excess of Alexa Fluor 488-C2 maleimide (Thermo Scientific), incubated for 5 mins and then flowed onto SMrTs. Images were acquired after 15 min of incubation followed by washing off the excess protein and dye. Membrane tube size in SMrTs was estimated using the supported bilayer formed in situ as a calibration standard (Dar *et al*., 2017). Fission index was estimated based on an image analysis routine. Images of tubes in SMrTs after they were exposed to Vps1 were background corrected using the mode value. A 512x512 pixel mask was generated using default threshold settings and a count of particles of area ≥ 1 pixel^2^ and circularity between 0-1 was estimated within this mask. This provides the total number of tube fragments. The same procedure was carried out for a smaller 20x512 pixel mask in the same image, which provides the total number of tubes. Fission index was calculated by dividing the number of tube fragments by the number of tubes and subtracting by 1. Fluorescence images were analyzed and rendered using Fiji. All data were plotted and analyzed using GraphPad Prism (version 10.0).

### Yeast strain generation

Gene deletion, replacement and tagging with fluorescent proteins were carried out according to Janke et al. (Janke *et al*., 2004). Primers were of HPLC-purified grade (Sigma). PCR amplicons generated using primers that are complementary to 50 bp upstream and downstream of the VPS1 locus were used to transform S288C cells according to Gietz et al. (Gietz *et al*., 1995). The *VPS1\** strain with mutants was generated in the *vps1Δ* background. Yeast strains used here are listed in Table S5.

### Yeast cell imaging and image analysis

Primary yeast cultures were grown on appropriate media overnight till saturation. Secondary cultures were started by diluting the primary culture to an OD of 0.1-0.2 and grown to an OD of 0.6-0.8. 1.5 ml of this culture was spun and resuspended in 0.2 ml of DPBS. 8-well Lab-Tek chambers (Thermo Fisher Scientific) were etched with 5 N NaOH for 30 min and washed with deionized water. Concanavalin A (ConA) (Sigma) at 0.5 mg/ml in DPBS was added to the chamber and incubated for 30 min. Chambers were washed with DPBS. 0.2 ml of the yeast cell suspension was incubated with the ConA-coated surface for 30 min, washed with DPBS 3-5 times and imaged in DPBS. For quantitative analysis of vacuolar localization, 512x512 pixel images of several fields of cells expressing cargo and the vacuolar marker Vph1-mSc were acquired. Images of cargo and Vph1-mSc were stacked separately and background corrected using the mode value. A mask was generated using the Vph1-mSc fluorescence with default threshold settings. The average fluorescence intensity of cargo and Vph1-mSc in particles of area ≥ 10 pixel^2^ and circularity between 0-1 was estimated within this mask. Vacuolar localization of cargo was depicted as a ratio of cargo to Vph1-mSc fluorescence. Normalization of cargo signals to Vph1-mSc was found to significantly correct for heterogeneity from distorted vacuole morphology, especially in *vps1Δ* cells, out-of-focus signals and variations in imaging conditions across experiments. Live cell imaging movies were bleach corrected using the ratio method applied through the bleach correction plugin in Fiji. For coefficient-of-variation (COV) analysis of ALFA-NB-mNG fluorescence, images were first corrected for background using the minima value. A mask was generated using default threshold settings from the ALFA-NB-mNG images. The mean and standard deviation of ALFA-NB-mNG fluorescence in particles of area ≥ 400 pixel^2^ and circularity between 0-1 was estimated. COV is expressed as the standard deviation divided by the average fluorescence. Fluorescence images were analyzed and rendered using Fiji.

### Spot growth assay

Cells were grown to an OD of 0.6-0.7, pelleted and resuspended in deionized water. Cultures were sequentially diluted 10-fold each time for 5 times in a 96 well plate. 5 μl of the culture was spotted on YPD agar. Cells were grown at 30 °C and 37 °C for 2 days and imaged.

### Western blotting

Cells were grown to an OD of 0.6-0.8, diluted to the exact same OD across strains and pelleted at 3200g for 5 min at 4 °C. The pellet was resuspended in 0.8 ml of cold autoclaved deionized water and immediately mixed with 0.15 ml of 1.85 N NaOH. Samples were vortexed and kept on ice for 5 min. 0.15 ml of 55 % (w/w) Trichloroacetic acid (TCA) was added to the samples, vortexed and kept on ice for 10 min. Samples were spun at 18,000 g for 20 min at 4°C and the supernatant was discarded. Samples were spun again at 18,000 g for 5 min to remove residual TCA. The pellet was resuspended in HU-DTT buffer (8 M urea, 5% SDS, 200 mM Tris-HCl pH 6.8, 0.1 mM EDTA, bromophenol blue,10 mM DTT) and heated at 99 °C for 20 min. Equal volumes of samples were loaded and resolved on a 10% SDS-PAGE gel and transferred to PVDF membrane. Blocking was done using 5% skimmed milk in 1X TBST (1X Tris Buffer Saline, 0.1% Tween 20). Blots were incubated with anti-ALFA-HRP antibody (NanoTag Biotechnologies) for 3 hrs in 5% skimmed milk and rinsed thrice with 1X TBST and imaged for chemiluminescence on an iBright 1500 (Invitrogen). Fluorescence images were rendered using Fiji.

### Sequence alignment

The *S. cerevisiae* InsB sequence was checked for homology with other fungal sequences using the Fungal Blast algorithm on the Saccharomyces Genome Database (Engel *et al*., 2025). Sequences showing significant similarities were collated and run through Clustal Omega (Madeira *et al*., 2024). Results in FASTA format converted by Seqret were rendered using SnapGene.

### Vacuole isolation

Yeast vacuoles were isolated according to previously described protocols (Cabrera and Ungermann, 2008). Cells were grown to an OD of 0.6-0.8 and pelleted at 3200 g for 5 min. Cell pellets were resuspended in DTT solution (0.1 mM Tris, pH 9.4, 10 mM DTT) and incubated in a water bath at 30 °C for 20 min. Cells were then pelleted again at 3200 g for 5 min and the supernatant was discarded. Cells were resuspended in spheroplasting buffer (0.1X YPD, 0.6 M sorbitol, 50 mM potassium phosphate, pH 7.5) with 1 mg/ml of Lyticase (Sigma, L4025-25KU) and incubated in a water bath at 30°C for 30 min.

Spheroplasts were pelleted at 5300 g for 5 min and resuspended in 15 % Ficoll (w/v) in PS buffer (10 mM Pipes/KOH, pH 6.8 with 200 mM sorbitol). This suspension was mixed with 0.4 mg/ml DEAE dextran (Sigma, 93556) and transferred to a Beckman Coulter SW 40 tube. The mixture was incubated on ice for 5 min and then shifted to 30 °C for 1.5 min. A step gradient of 8%, 4%, and 0% Ficoll (w/v) in PS buffer was then layered on top of the mixture. Tubes were spun at 1,20,000 g for 90 min at 4°C in a SW40 rotor. The vacuole fraction at the interface of 0% and 4% Ficoll was carefully collected. The vacuole fraction was solubilized using 1% Triton X-100 and protein concentration was measured with BCA using BSA as standard.

### Proteomic analysis

Vacuole samples isolated from *vps1Δ* and wildtype cells and containing 50 μg of total protein were separately precipitated using a methanol, chloroform and water mixture (Wessel and Flügge, 1984). Precipitates were resuspended in 100 mM TEAB buffer containing 8 M urea. 5 mM of DTT was added to the mixture and samples were incubated at 60 °C for 30 min. Samples were cooled down to room temperature and 15 mM of iodoacetamide was added to the mixture and incubated in the dark for 15 min. The solution was diluted to a urea concentration of 1M with TEAB buffer and incubated with 0.5 mg/ml of sequencing grade modified Trypsin (Promega, V5111) at 37 °C for 16 hr. Tryptic digests were processed for quantitative proteomics using reductive dimethylation (ReDi) labeling (Boersema *et al*., 2009). Tryptic peptides from *vps1Δ* and wildtype vacuoles were labeled with 4% heavy formaldehyde (CD_2_O) and 4% light formaldehyde (CH2O), respectively in 0.6 M sodium cyanoborohydride (NaBH3CN) and incubated at room temperature for 1 hr. Labelling reaction was quenched by adding 1% ammonium hydroxide (v/v) and acidified by adding 1% trifluoroacetic acid. The solution was spun at 18000 g for 20 min at room temperature to get rid of particulate matter and the supernatant was collected. Equal volumes of the supernatant from the heavy and light labelled samples were mixed and desalted twice using a C18 column. Samples were dried and stored at -80 °C. Samples were resuspended in 0.1% formic acid and analyzed through an Eksigent nanoLC 425 attached to a Sciex Triple-TOF 6600 mass spectrometer.

Samples were first passed through an Eksigent C18 trap column to get rid of any common contaminants and eluted using a linear acetonitrile gradient. Proteomics data acquisition was carried out on an information-dependent acquisition (IDA) mode over a specified mass range of 300-2000 m/z. Peptide identification and quantitation was done using Protein Pilot software. *S. cerevisiae* (S288C) protein database (Uniprot release date, 2021) was used as the peptide library. Data from 2 biological and 6 technical repeats were pooled and analyzed together. Iodoacetamide was specified as alkylating agent and dimethylation was specified as extra modification. Identification was performed on peptides filtered with <1% FDR. Proteins with ≥ 2 peptides and with a -log_10_(*p*-value) of 0.2 were considered.

## Data analysis

All data were plotted and analyzed using GraphPad Prism (version 10.0).

## Acknowledgements

S.G. and U.S. thank the Council of Scientific and Industrial Research (CSIR) for graduate student fellowships. G.S. thanks the Kishore Vaigyanik Protsahan Yojana (KVPY) and IISER Pune for undergraduate fellowships. T.J.P. thanks the Howard Hughes Medical Institute for an International Research Scholar’s Grant (Grant No. 55008746), the Anusandhan National Research Foundation for a SUPRA Grant (Grant No. SPR12021100014), and the DBT/Wellcome Trust India Alliance for a Team Science Grant (Grant No. IA/TSG/21/1/600245). We thank Saravanan Palani (Indian Institute of Science, India) for help with strains, reagents and protocols. We thank members of the Pucadyil lab for helpful comments on the manuscript.

## Author Contributions

S.G., G.S., U.S. and T.J.P. conceived of experiments and developed methods. S.G., G.S. and U.S. performed experiments. S.G., G.S. and T.J.P. analyzed data. S.G. wrote the first draft of the manuscript which worked on by T.J.P. S.G., G.S., U.S. and T.J.P reviewed and edited the manuscript. T.J.P. acquired financial support for this study.

## Supplementary Tables

**Table S1.** List of proteins identified in the quantitative proteomics analysis.

**Table S2.** Identified proteins sorted based on cellular component annotation using the GO slim mapping tool in the Saccharomyces Genome Database.

**Table S3.** Depleted proteins sorted based on cellular process annotation using the GO slim mapping tool in the Saccharomyces Genome Database.

**Table S4.** Plasmids used in this study.

**Table S5.** Yeast strains used or generated in this study.

## Supplementary Movies

**Movie S1. Vps1-catalyzed membrane fission.** Time lapse movie showing the effect of flowing Vps1 mixed with GTP on PC:PS:PI(3)P templates.

**Movie S2. Vps1-catalyzed membrane fission depends on substrate size.** Time lapse movie showing the effect of flowing Vps1 mixed with GTP on PC:PS:PI(3)P tubes of ∼15 nm and ∼40 nm radius.

**Movie S3. Tube constriction during Vps1-catalyzed membrane fission.** Time lapse movie showing the effect of flowing GTP on preassembled GppNHP-bound Vps1 scaffolds on a tube.

**Movie S4. Vps10 dynamics in wildtype cells.** Time lapse movie showing the distribution and dynamics of Vps10-mNG and Vph1-mSc in wildtype cells.

**Movie S5. Vps10 dynamics in *vps1****Δ* **cells.** Time lapse movie showing the distribution and dynamics of Vps10-mNG and Vph1-mSc in *vps1Δ* cells.

**Movie S6. Vps10 dynamics in Vps1* cells.** Time lapse movie showing the distribution and dynamics of Vps10-mNG and Vph1-mSc in *Vps1\** cells.

**Movie S7. Vps1 dynamics in cells.** Time lapse movie showing the distribution and dynamics of ALFA- NB-mNG in *VPS1\** cells.

**Movie S8. Vps1(K42A) dynamics in cells.** Time lapse movie showing the distribution and dynamics of ALFA-NB-mNG in *VPS1*(K42A) cells*.

**Movie S9. Vps1(G436D) dynamics in cells.** Time lapse movie showing the distribution and dynamics of ALFA-NB-mNG in *VPS1*(G436D)*.

**Movie S10. Vps1(Δ4F) dynamics in cells.** Time lapse movie showing the distribution and dynamics of ALFA-NB-mNG in *VPS1*(Δ4F)*.

**Movie S11. Vps1(Δ3K) dynamics in cells.** Time lapse movie showing the distribution and dynamics of ALFA-NB-mNG in *VPS1*(Δ3K)*.

**Movie S12. Vps1(ΔInsB) dynamics in cells.** Time lapse movie showing the distribution and dynamics of ALFA-NB-mNG in *VPS1*(ΔInsB)*.

**Movie S13. Neo1 dynamics in wildtype cells.** Time lapse movie showing the distribution and dynamics of Neo1-mNG and Vph1-mSc in wildtype cells.

**Movie S14. Neo1 dynamics in *vps1****Δ* **cells.** Time lapse movie showing the distribution and dynamics of Neo1-mNG and Vph1-mSc in *vps1Δ* cells.

**Movie S15. Lem3 dynamics in wildtype cells.** Time lapse movie showing the distribution and dynamics of Lem3-mNG and Vph1-mSc in wildtype cells.

**Movie S16. Lem3 dynamics in *vps1****Δ* **cells.** Time lapse movie showing the distribution and dynamics of Lem3-mNG and Vph1-mSc in *vps1Δ* cells.

---

**Fig. S1.**
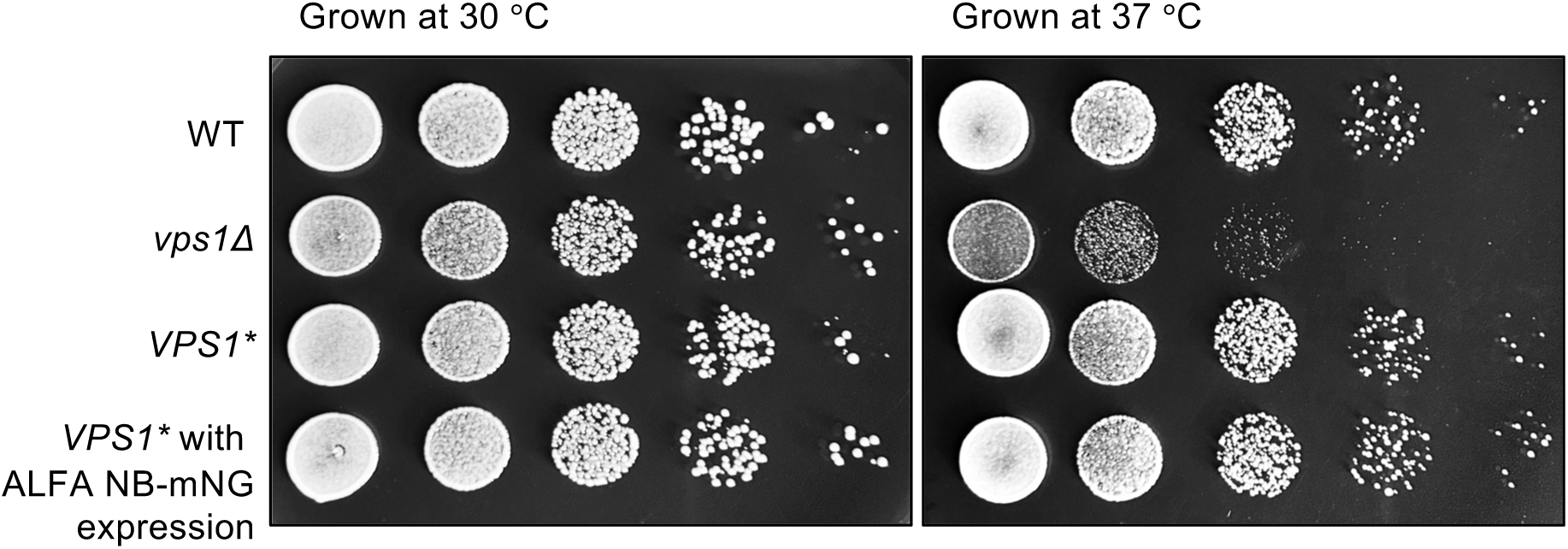
Growth characteristics of *VPS1\** strains. Spot assay comparing growth of WT, *vps1Δ* and *VPS1\** alone or expressing the ALFA-NB-mNG at 30 °C and 37 °C.

**Fig. S2.**
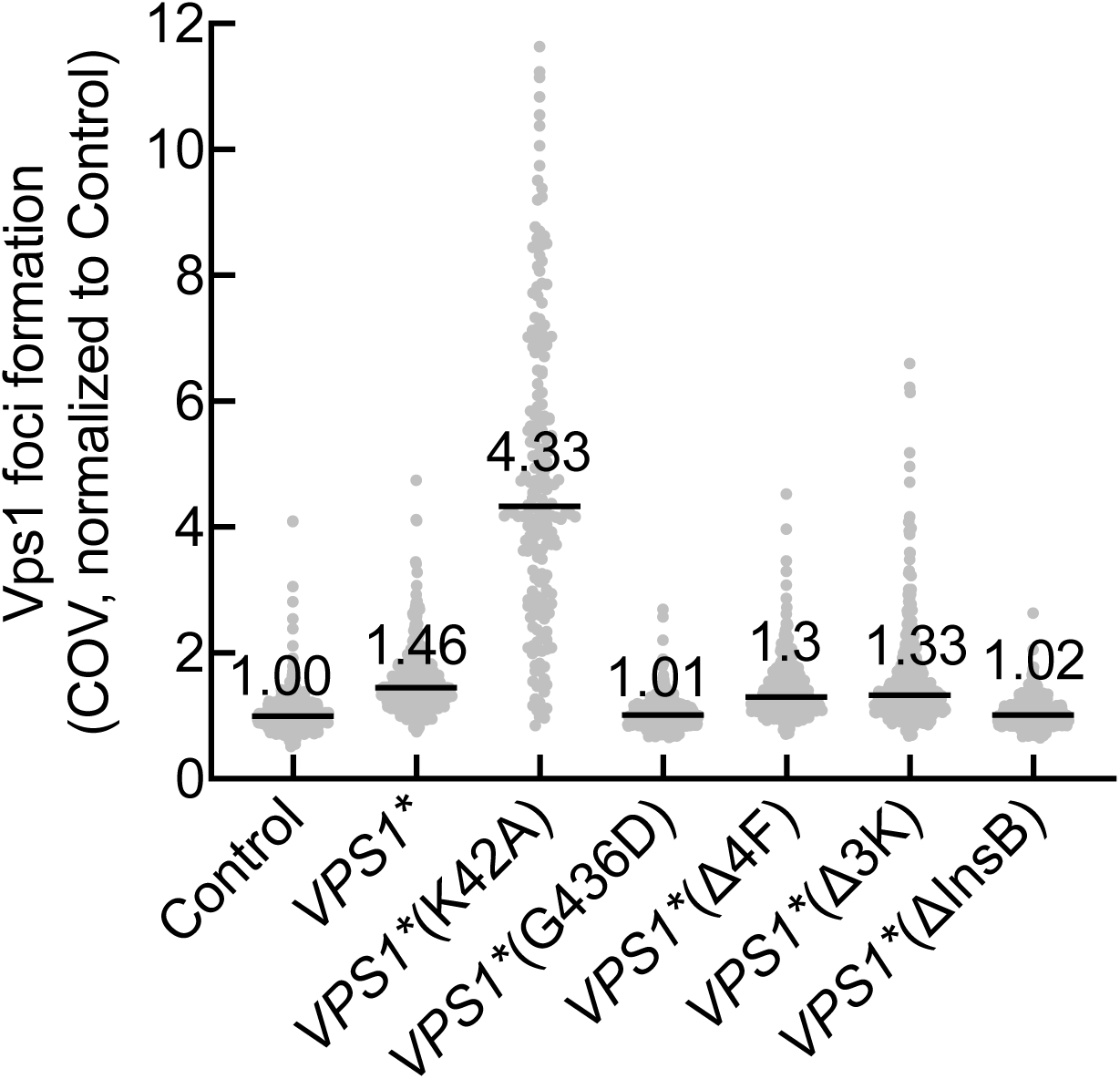
Quantitation of foci-like distribution of Vps1*. Plot showing the COV of ALFA-NB-mNG fluorescence in 563 cells not expressing *VPS1\** (Control), 398 for *VPS1*,* 197 for *VPS1*(K42A)*, 370 for *VPS1*(G436D)*, 403 for *VPS1*(Δ4F)*, 421 for *VPS1*(Δ3K)*, and 509 for *VPS1*(ΔInsB)*. Number and black lines represent median value. Data are normalized to the median seen in Control cells.

**Fig. S3.**
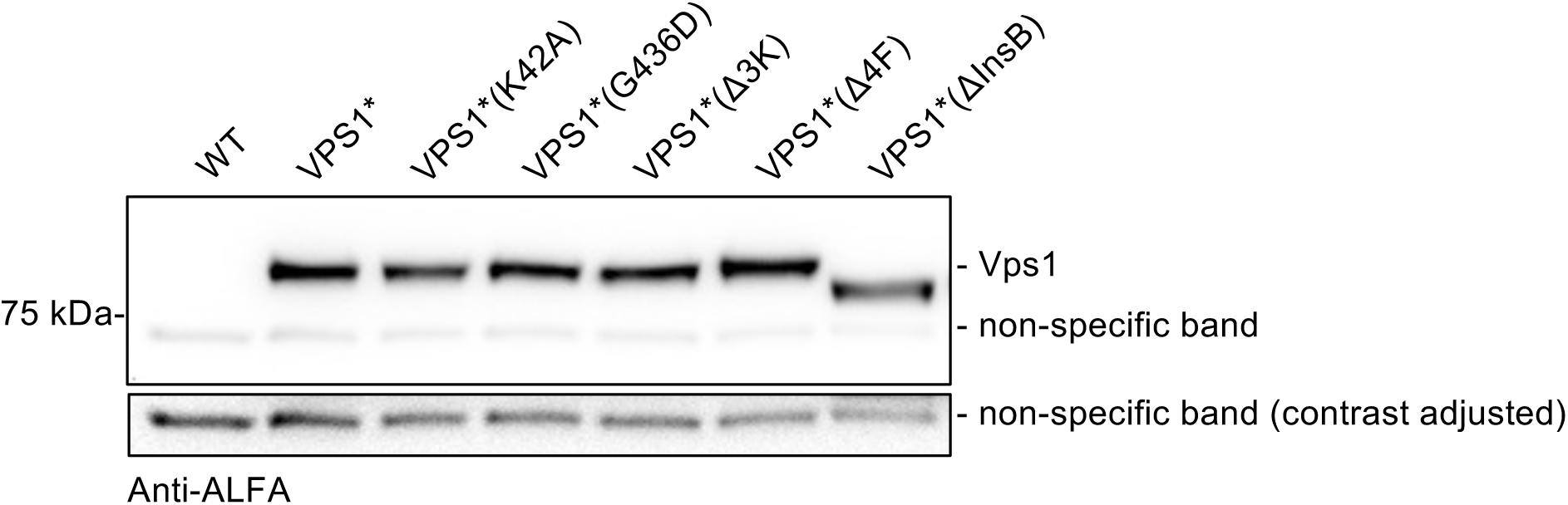
Western blots of Vps1* and mutants. Lysates were blotted with the ALFA antibody, which also detects a non-specific band at ∼75 kDa that served as loading control (Akhuli *et al*., 2022).

## Notes

### Competing Interest Statement

The authors have declared no competing interest.

